# Optical microscopy reveals the dynamic nature of *B. pseudomallei* morphology during β-lactam antimicrobial susceptibility testing

**DOI:** 10.1101/2020.01.13.904995

**Authors:** Heather P. McLaughlin, Julia Bugrysheva, David Sue

**Affiliations:** Laboratory of Preparedness and Response Branch, Division of Preparedness and Emerging Infections, National Center for Emerging and Zoonotic Infectious Diseases, Centers for Disease Control and Prevention, 1600 Clifton Road NE, MS: H17-5, Atlanta, GA 30333, USA

**Keywords:** Cell morphology, *B. pseudomallei*, β-lactam antibiotics

## Abstract

**Background:** In Gram-negative species, β-lactam antibiotics target penicillin binding proteins (PBPs) resulting in morphological alterations of bacterial cells. Observations of antibiotic-induced cell morphology changes can rapidly and accurately differentiate drug susceptible from resistant bacterial strains; however, resistant cells do not always remain unchanged. *Burkholderia pseudomallei* is a Gram-negative, biothreat pathogen and the causative agent of melioidosis, an often fatal infectious disease for humans.

**Results:** Here, we identified β-lactam targets in *B. pseudomallei* by *in silico* analysis. Ten genes encoding putative PBPs, including PBP-1, PBP-2, PBP-3 and PBP-6, were detected in the genomes of susceptible and resistant strains. Real-time, live-cell imaging of *B. pseudomallei* strains demonstrated dynamic morphological changes in broth containing clinically relevant β-lactam antibiotics. At sub-inhibitory concentrations of ceftazidime (CAZ), amoxicillin-clavulanic acid (AMC), and imipenem (IPM), filamentation, varying in length and proportion, was an initial response of the multidrug-resistant strain Bp1651 in exponential phase. However, a dominant morphotype reemerged during stationary phase that resembled cells unexposed to antibiotics. Similar morphology dynamics were observed for AMC-resistant strains, MSHR1655 and 724644, when exposed to sub-inhibitory concentrations of AMC. For all *B. pseudomallei* strains evaluated, increased exposure time and exposure to increased concentrations of AMC at and above minimal inhibitory concentrations (MICs) in broth resulted in cell morphology shifts from filaments to spheroplasts and/or cell lysis. *B. pseudomallei* morphology changes were more consistent in IPM. Spheroplast formation followed by cell lysis was observed for all strains in broth containing IPM at concentrations greater than or equal to MICs, however, the time to cell lysis was variable. Length of *B. pseudomallei* cells was strain-, drug- and drug concentration-dependent.

**Conclusions:** Both resistant and susceptible *B. pseudomallei* strains exhibited filamentation during early exposure to AMC and CAZ at concentrations used to interpret susceptibility (based on CLSI guidelines). While developing a rapid β-lactam antimicrobial susceptibility test based on cell-shape alone requires more extensive analyses, optical microscopy detected *B. pseudomallei* growth attributes that lend insight into antibiotic response and antibacterial mechanisms of action.

## Background

The World Health Organization (WHO) identified antimicrobial resistance as one of the most important problems for human health that threatens the effective prevention and treatment of infectious diseases [1]. A 2019 Centers for Disease Control and Prevention (CDC) report on antibiotic resistance threats highlights the latest burden estimates for human health in the U.S., listing 18 resistant pathogens into one of three categories: urgent, serious and concerning [2]. Timely administration of appropriate drug therapy is essential for both patient outcomes and for combatting the spread of antibiotic resistance [3, 4]. β-lactams are the most common treatment for bacterial infections and the class accounts for 70% of antibiotic prescriptions in the United States [5]. However, increased exposure of bacteria to a multitude of β-lactams drives adaptation and has led to the production and mutation of β-lactamases, resulting in resistance [6].

Melioidosis is a life-threatening human infection with case fatality rates that may exceed 70% as a result of ineffective antimicrobial therapy [7-9]. Naturally-acquired melioidosis infections are caused by inhalation, ingestion or exposure of broken skin to the pathogen *Burkholderia pseudomallei*. This disease is endemic across tropical areas and is estimated to account for ∼165,000 human cases per year worldwide, ∼89,000 of which result in death [8]. The United States Federal Select Agent Program includes *B. pseudomallei* as a Tier 1 biological select agent. Public health and safety could be compromised if this pathogen was deliberately misused due to ease of propagation, small infectious dose, and high mortality rate. Awareness of melioidosis and research into *B. pseudomallei* is increasing due to the heavy disease burden and the biothreat potential [10].

β-lactams ceftazidime (CAZ), amoxicillin-clavulanic acid (AMC), and imipenem (IPM) are antibiotics used for melioidosis treatment [9]; however, treatment failures have been attributed to acquired and intrinsic *B. pseudomallei* drug resistance [11-13]. Drug inactivation and drug target modification are described as mechanisms of resistance. Mutations resulting in amino acid changes in and upstream of the β-lactamase gene *penA* confer resistance to AMC, CAZ and IPM in strain Bp1651 and to AMC in strain MSHR1655 [14-16]. In addition, a reversible gene duplication and amplification event in a chromosomal region containing *penA* resulted in acquired CAZ resistance [17]. Loss of the drug target penicillin-binding protein 3 (PBP-3) also contributes to CAZ-resistance in *B. pseudomallei* [18].

Inactivation of specific PBPs targeted by β-lactam antibiotics induces well-defined morphological changes in other Gram negative bacteria: (i) inhibition of PBP-3 leads to formation of filaments, (ii) inhibition of PBP-2 results in the production of round cells and cell-wall deficient spheroplasts, (iii) and inhibition of PBP-1A and PBP-1B induces rapid cell lysis [19]. To date, three PBP-3 homologs have been reported in *B. pseudomallei* [18], but genes encoding putative PBP-2 and PBP-1 have not been identified. Cell morphology can manifest differently when β-lactams demonstrate an affinity for multiple PBP targets. Moreover, morphological response is dependent on the number of target PBPs present, the antibiotic concentration, and the specificities of enzyme binding sites [19]. Documenting β-lactam-induced morphology changes could improve our understanding of antibiotic response and mechanisms of action as well as help identify trends that are predictive of *B. pseudomallei* susceptibility.

Early administration of effective drug therapy is critical for positive melioidosis patient outcomes. Rapid phenotypic β-lactam antimicrobial susceptibility testing (AST) of *B. pseudomallei* can facilitate the administration of antibiotics with confirmed activity against infecting strains. Optical screening was previously used for the rapid AST of Gram negative biothreat agents to several classes of antibiotics including aminoglycosides, tetracyclines, fluoroquinolones and β-lactams [20]. The time required to accurately determine susceptibility decreased by up to 70% compared to conventional broth microdilution (BMD) testing, but most of the antibiotic agents tested did not induce cell filamentation. Microbial growth was measured by estimating bacterial cell surface area and the automated assay could not differentiate between cell elongation and cell division of β-lactam-induced filamentous cells, including some *B. pseudomallei* strains grown in the presence of CAZ [20]. However, real-time video imaging by microscopy revealed antibiotic-induced cell morphology changes.

Assessment of cell morphology, rather than bacterial density, was previously used to rapidly differentiate susceptible and resistant Gram negative bacteria in the presence of β-lactams [21-23] and rapid AST based on these analyses showed high categorical agreement compared to gold standard BMD results. The morphological response of resistant strains was variable between studies; some reported that cell shape remained unchanged in the presence of β-lactams, while others observed cell swelling and filament formation. Here, we use optical microscopy to i) study morphological responses of drug resistant and susceptible *B. pseudomallei* strains in broth containing β-lactams, ii) quantify cell length, shape, and response during exposure to β-lactams, and iii) investigate the usefulness of cell morphology for rapid susceptibility testing of *B. pseudomallei* to β-lactams. We describe the utility of optical screening to explore trends in morphology and capture growth characteristics that maybe be indicative of specific antimicrobial response. We also identify PBP homologs encoded in the *B. pseudomallei* genome which may represent the potential targets for β-lactams antibiotics and better explicate the antibacterial mechanisms of action.

## Materials and Methods

### Bacterial strains, growth conditions and biosafety procedures

Seven *B. pseudomallei* strains, including one multidrug-resistant (MDR) and two AMC-resistant (AMC-R) strains, were included in this study (**Tables 1-3).** Bacterial stocks were maintained in 20% glycerol at −70°C and cultured on trypticase soy agar II with 5% sheep’s blood (SBA) (Fisher Scientific, Pittsburg, PA) at 35°C in ambient air for testing. All work with *B. pseudomallei* was completed inside a class II type A2 biological safety cabinet located in a BSL-3 laboratory registered with the U.S. Federal Select Agent Program and is subject to select agent regulations (42-CFR-Part-73). Procedures were performed by trained personnel wearing a powered air-purifying respirator and protective laboratory clothing [24].

**Table 1.** Antimicrobial susceptibility profile and morphological analysis of MDR Bp1651.

| MIC Interpretative Criteria (CLSI) |  |  |  | Strain | Filamentation (4h) |  |  |  | Time (h) cell lysis begins |
| --- | --- | --- | --- | --- | --- | --- | --- | --- | --- |
| β-lactam (μg/ml) | S | I | R | Bp1651 MIC | $\frac{1}{2}$ MIC | | MIC | | MIC |
| | | | | | # Cells $\geq 15 \mu$m | Median CL (μm) | # Cells $\geq 15 \mu$m | Median CL (μm) | |
| AMC | $\leq 8/4$ | 16/8 | $\geq 32/16$ | 64/32 (R) | 6/100 | 5 | 0/100 | 4 | $3.7 \pm 0.0$ |
| CAZ | $\leq 8$ | 16 | $\geq 32$ | >128(R) | 49/100* | 14* | 39/100* | 10.5* | filaments and spheroplasts* |
| IPM | $\leq 4$ | 8 | $\geq 16$ | 32 (R) | 8/100 | 4 | 0/100 | 3 | $3.6 \pm 0.2$ |

Minimal inhibitory concentrations (MIC) of amoxicillin-clavulanic acid (AMC), ceftazidime (CAZ), and imipenem (IPM) were determined by conventional BMD based on CLSI guidelines. Susceptibility interpretation is defined as resistant (R), intermediate (I) or susceptible (S). Cell length (CL) was measured after 4 h and median cell length was calculated (n=100 cells). (\*) filamentation and cell lysis data in the presence of CAZ was calculated using a value of 128 μg/ml for the MIC due to concentration range in Sensititre drug panel for testing. Time to cell lysis is represented as the mean ± SD of triplicate samples based on growth kinetic data and visual observation of video imaging.

**Table 2.** Morphological analysis of *B. pseudomallei* strains in the presence of AMC and CAZ.

| Strain | AMC<br>MIC<br>($\mu\text{g/ml}$) | Filamentation (4h) | | | | Time (h) cell lysis begins | | |
| --- | --- | --- | --- | --- | --- | --- | --- | --- |
| | | $\frac{1}{2}$ MIC | | MIC | | MIC | 2x MIC | 4x MIC |
| | | # Cells<br>$\geq 15 \mu\text{m}$ | Median<br>CL ($\mu\text{m}$) | # Cells<br>$\geq 15 \mu\text{m}$ | Median<br>CL ($\mu\text{m}$) | | | |
| MSHR1655 | 32/16 (R) | 62/100 | 19.5 | 61/100 | 19 | $6.1 \pm 0.2$ | $4.0 \pm 0.3$ | not tested |
| 724644 | 16/8 (I) | 93/100 | 59 | 24/100 | 6 | $4.8 \pm 1.0$ | $2.9 \pm 0.2$ | $2.9 \pm 0.2$ |
| 6788 | 8/4 (S) | 86/97 | 43 | 82/100 | 42.5 | remains<br>filamentous | filaments and<br>spheroplasts | only<br>spheroplasts |
| 1026b | 4/2 (S) | 42/100 | 12 | 80/97 | 44 | remains<br>filamentous | remains<br>filamentous | $5.2 \pm 0.4$ |

| Strain | CAZ<br>MIC<br>($\mu\text{g/ml}$) | Filamentation (4h) | | | | Time (h) cell lysis begins | | |
| --- | --- | --- | --- | --- | --- | --- | --- | --- |
| | | $\frac{1}{2}$ MIC | | MIC | | MIC | 2x MIC | 4x MIC |
| | | # Cells<br>$\geq 15 \mu\text{m}$ | Median<br>CL ($\mu\text{m}$) | # Cells<br>$\geq 15 \mu\text{m}$ | Median<br>CL ($\mu\text{m}$) | | | |
| 724644 | 8 (S) | 93/100 | 56.5 | 95/100 | 59.5 | remains<br>filamentous | remains<br>filamentous | remains<br>filamentous |
| MSHR1655 | 4 (S) | 48/100 | 13 | 61/100 | 22 | $8.2 \pm 0.4$ | $8.0 \pm 0.0$ | $8.0 \pm 0.3$ |
| 1026b | 2 (S) | 28/100 | 8 | 94/100 | 36 | remains<br>filamentous | remains<br>filamentous | remains<br>filamentous |

The minimal inhibitory concentrations (MIC) of amoxicillin-clavulanic acid (AMC) and ceftazidime (CAZ) were determined by conventional BMD testing based on CLSI guidelines. Susceptibility interpretations are defined as resistant (R), intermediate (I), or susceptible (S). Cell length (CL) was measured at time 4 h and median cell length at MICs was calculated (n=95-100 cells). Time to cell lysis is represented as the mean ± SD of triplicate samples based on growth kinetic data and visual observation of video imaging. Blue indicates breakpoint for susceptibility.

**Table 3.** Optical screening-based IPM susceptibility testing and morphological analysis of susceptible *B. pseudomallei*.

| Strain | IPM<br>MIC<br>(µg/ml) | Filamentation<br>(# Cells ≥15 µm)<br>½ MIC | Time (h) to determine<br>IPM susceptibility |  | Time (h) cell lysis begins |  |  |
| --- | --- | --- | --- | --- | --- | --- | --- |
|  |  |  | 8 µg/ml | 16 µg/ml | MIC | 2x MIC | 8 µg/ml |
| 724644 | 2 | no | 1.7 ± 0.0 | 1.7 ± 0.0 | 3.0 ± 0.0 | 2.9 ± 0.2 | 3.0 ± 0.0 |
| MSHR1655 | 1 | yes (15/100) | 1.3 ± 0.9 | 1.7 ± 0.5 | 3.9 ± 0.4 | 4.0 ± 0.7 | 3.6 ± 0.2 |
| 1026b | 1 | no | 3.8 ± 0.7 | 2.2 ± 1.7 | 5.0 ± 0.7 | 5.3 ± 0.6 | 5.7 ± 0.9 |
| 1631 | 0.5 | no | 3.7 ± 0.5 | 3.7 ± 0.0 | 9.1 ± 0.5 | 8.2 ± 0.8 | 4.6 ± 0.2 |
| 6296 | 0.25 | yes (6/100) | 3.7 ± 0.0 | 3.3 ± 0.5 | 9.8 ± 0.2 | 8.3 ± 1.4 | 5.3 ± 0.3 |

The minimal inhibitory concentration (MIC) of imipenem (IPM), determined by conventional BMD, and the time (h) required to determine susceptibility of *B. pseudomallei* strains (8 µg/ml and 16 µg/ml IPM) with a confidence level (CL) of 95% ( $p \leq 0.05$ ) are listed. All strains were monitored by optical screening 18-20 h in the presence of IPM below, at and above MICs. Cell length was measured after 4 h. Detection of filamentation in sub-inhibitory concentrations and cell lysis at inhibitory concentrations was achieved by analysis of growth kinetic data and visual observation of real-time video imaging.

### Antimicrobials and susceptibility testing by BMD

Antimicrobial susceptibility profiles for each strain are listed in **Tables 1-3**. Minimal inhibitory concentrations (MIC) were first determined by conventional BMD testing following CLSI guidelines for medium, inoculum, and incubation temperature [25]. BMD susceptibility testing panels were prepared with Cation-Adjusted Mueller Hinton Broth (CAMHB) in house. β-lactam antibiotics selected for this study were amoxicillin-clavulanic acid (AMC) (Toku-E, Bellingham, WA and USP, Frederick, MD), ceftazidime (CAZ) (Sigma Aldrich, St. Louis, MO) and imipenem (IPM) (Toku-E, Bellingham, WA). Two-fold antibiotic concentrations were tested ranging from 0.06/0.03-128/64 µg/ml AMC, 0.06-128 µg/ml CAZ, and 0.03-64 µg/ml IPM. *B. pseudomallei* strains were classified as resistant (R), intermediate (I), or susceptible (S) based on interpretive criteria outlined by CLSI [25]. MICs were recorded after 16 to 20 h of incubation at 35°C ambient air, with the exception of MSHR1655 (43 h). Incubation was extended due to insufficient growth in the control well of the BMD panel.

### Susceptibility testing by optical screening

An optical screening instrument, the oCelloScope (BioSense Solutions ApS, Farum, Denmark), was used for rapid β-lactam AST of *B. pseudomallei* strains. As previously described in McLaughlin *et al*. [20], 96-well Sensititre panels (Trek Diagnostics, ThermoFisher Scientific) containing desiccated AMC, CAZ and IPM were inoculated with *B. pseudomallei* cell suspensions in CAMHB with N-tris (hydroxymethyl) methyl-2-aminoethanesulfonic acid (TES) (Remel Inc., Lenexa, KS). From overnight SBA culture growth, inocula were prepared by making a cell suspension in CAMHB to a turbidity equivalent to 0.5 McFarland standard followed by a 1:50 dilution in CAMHB. Antibiotic concentrations evaluated by optical screening-based susceptibility testing were 2/1-64/32 µg/ml AMC, 1-128 µg/ml CAZ, and 0.12-32 µg/ml IPM. For drug concentrations tested below the Sensititre panel range, 1:2 dilutions of the desiccated antibiotic were made using the inocula as the diluent. Cell suspensions (90 µl) were transferred to a 96-well flat bottom plate, sealed with a breathable film cover (Breathe-Easy Sealing Membranes, Sigma Aldrich, St. Louis, MO) and monitored over time by optical screening at 35°C in ambient air. Instrument-derived growth values were recorded every 20 minutes for 18 to 24 h. For each strain, susceptibility testing was performed with three technical replicates in two biological experiments.

### Growth kinetics and data analysis

Automated growth kinetic experiments were performed using the Segmentation and Extraction Surface Area (SESA) algorithm of the oCelloScope-specific software, UniExplorer (v. 5.0.3). Using this algorithm, bacteria were identified in a scan area based on contrast against the background, and growth values were calculated by summarizing bacterial surface area. Growth kinetic graphs represent the average growth values (n=3) ± standard deviations (SD), where indicated, from one representative experiment. Statistical analysis of growth data defined the time (h) required to determine antimicrobial susceptibility. The statistical significance, with a confidence level of 95% (*p*-value < 0.05), between a susceptible strain grown in media with and without β-lactam antibiotics over time was calculated using a two-tailed t-test. The minimum incubation times required for β-lactam AST are reported as the average ± SD from duplicate biological experiments (n=3). Cell lysis was observed by video imaging and the time (in h) was recorded when growth values began to decrease continuously over time. Cell lysis time is represented as the mean time (n=3) ± SD.

### Cell morphology imaging by optical screening

Real-time imaging of *B. pseudomallei* broth cultures in microtiter panel wells was performed simultaneously with β-lactam AST using the oCelloScope instrument. Images captured in CAMHB media without drug and in media containing either AMC, CAZ, or IPM at final concentrations below, at and above MICs during exponential and stationary phase growth. From each well containing 90 µl of culture, 10 images were taken from a tilted image plane to produce a z-stack image and videos were composed of z-stack images acquired over time. A ‘best focused’ image of 10 images was automatically selected through the UniExplorer Software and is depicted in figures. Times (h) in which strains were exposed to antibiotics are indicated in each figure legend.

### Analysis of bacterial cell length

The Segmentation task of UniExplorer was used to analyze individual cells from optical screen images of *B. pseudomallei*. Unless otherwise indicated, cell length was measured after 4 h utilizing the ‘thinned length’ object feature. For each strain analyzed, 95 to 100 cells were measured in media with and without the presence of antibiotic and the distribution of cell lengths were graphically represented by histogram or dot plot. The median cell length value is the midpoint of this distribution.

### Identification of penicillin binding protein homologs

Putative PBPs in *B. pseudomallei* 1026b were identified using the search feature of the UniProtKB database (https://www.uniprot.org/uniprot/), an online resource for protein sequence and annotation data. The Pfam database (http://pfam.xfam.org/) was utilized to predict conserved protein domains. Genes encoding putative PBPs in the 1026b genome (NCBI accession # CP002833, CP002834) were used as queries to identify corresponding homologs in Bp1651 (NCBI accession # CP012041, CP012042), MSHR1655 (NCBI accession # CP008779, CP008780), and Bp6296 (NCBI accession # CP018393, CP018394), in which completed assembled genomes are publicly available on NCBI. The nearest PBP protein homologs in *Pseudomonas aeruginosa* PAO1 (NCBI accession # NP 253108) were found using NCBI Protein BLAST and percent identities were based on amino acid sequence alignments.

## Results

### β-lactam-induced cell morphology dynamics of the multidrug-resistant strain Bp1651

*B. pseudomallei* strain Bp1651 is resistant to AMC, CAZ, and IPM based on CLSI MIC interpretive criteria (**Table 1**). Growth in broth culture of this MDR strain was monitored in real-time by optical screening in the presence of each β-lactam and in CAMHB only. AMC, CAZ, and IPM concentrations tested included the CLSI breakpoint for susceptibility and the three successive two-fold higher concentrations. Optical screening images captured Bp1651 cell morphology during exponential and stationary phase growth (**Fig. 1A**). In media without antibiotics, cells of typical *B. pseudomallei* length (≤ 5 µm) were observed for Bp1651 throughout all phases of growth. In sub-inhibitory concentrations of AMC, CAZ and IPM, filamentation was detected during exponential phase (**Fig. 1A**).

**Figure 1.**
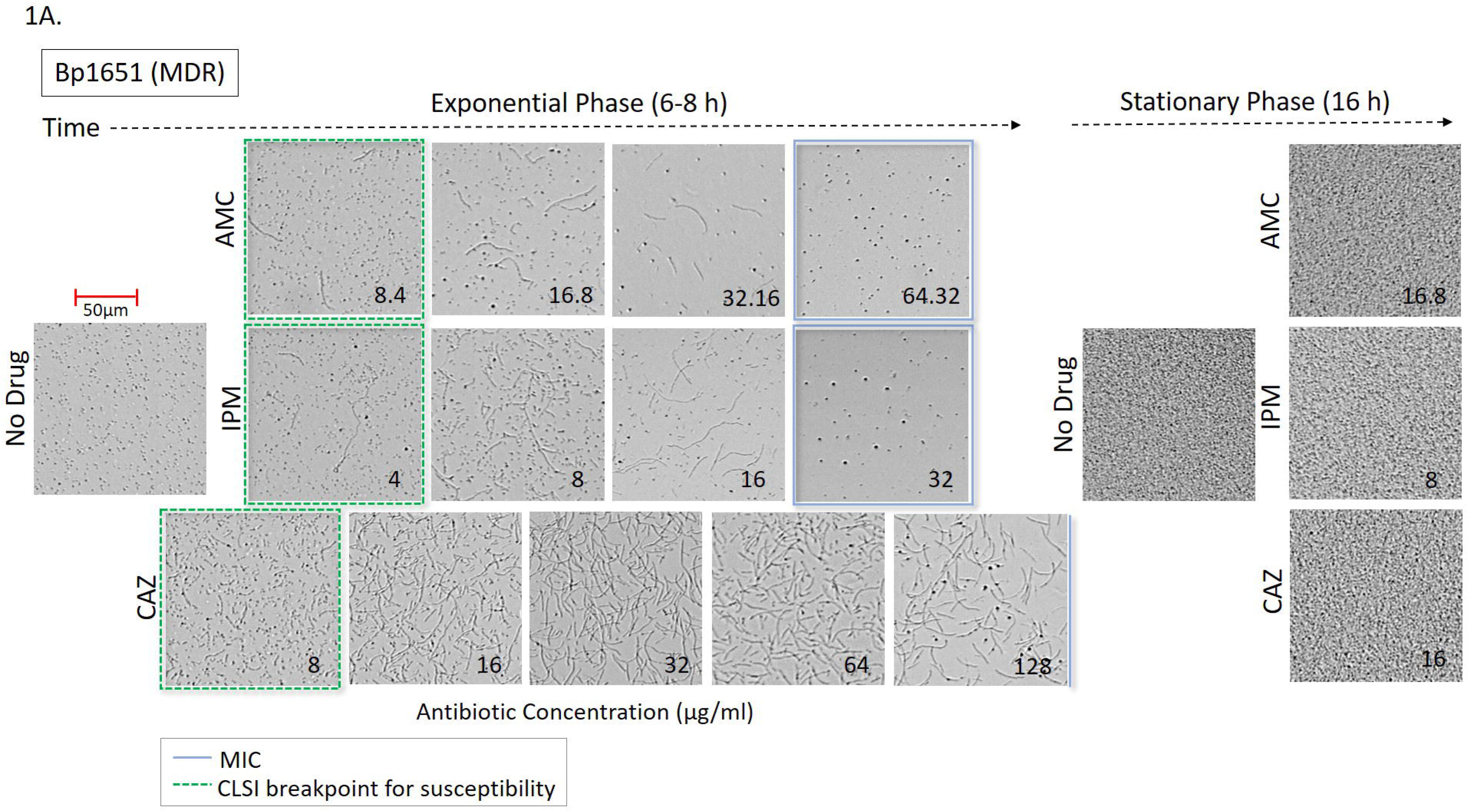
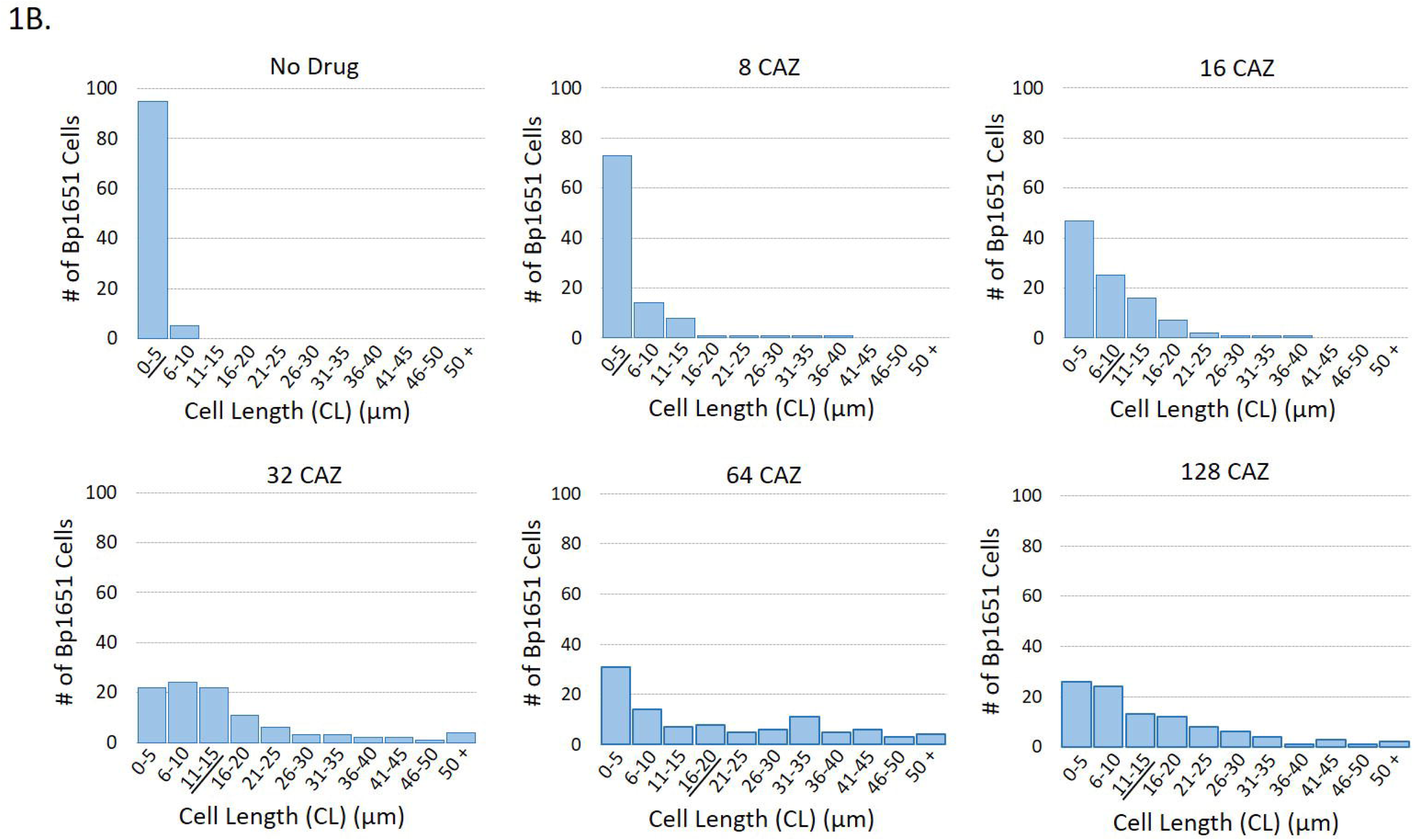
Cell morphology of the MDR Bp1651 strain cultured in the presence and absence of β-lactam antibiotics (AMC, CAZ, and IPM) (A). Optical screen images were captured during exponential and stationary phase growth. Drug concentrations (µg/ml) corresponding to the CLSI breakpoint for susceptibility (dotted green squares) and three or four successive two-fold increasing concentrations. Blue squares represent the Bp1651 MICs and the blue line indicates the CAZ MIC is ≥ 128 µg/ml. Distribution of cell lengths of Bp1651 cultured in the presence and absence of CAZ (µg/ml) (B). Histograms represent cells (n=100) measured (µm) after 4 h using the thinned length object feature. Multidrug-resistant (MDR), amoxicillin-clavulanic acid (AMC), ceftazidime (CAZ), and imipenem (IPM).

As outlined by CLSI, drug resistance or susceptibility of *B. pseudomallei* by BMD testing is assessed by observations of growth or inhibition of growth at the two-fold concentrations above and below 16/8 µg/ml AMC, 8 µg/ml IPM and 16 µg/ml CAZ. At these concentrations, the lengths of 100 Bp1651 cells were each measured after 4 h using the ‘thinned length’ object feature of the UniExplorer Software. Approximately 20% of cells exhibited filamentation (lengths ≥ 15 µm) in the presence of 16/8 µg/ml AMC. More than half of cells (62/100) remained ≤ 5 µm, similar to cells measured in broth alone. The longest cell recorded at this concentration was 29 µm (data not shown). After 4 h at 8 µg/ml IPM and 16 µg/ml CAZ, some filamentous cells were observed with the longest cells measured at 37 µm and 46 µm, respectively. About half of Bp1651 cells remained ≤ 5 µm at these concentrations of IPM (56/100 cells) and CAZ (47/100 cells). Bp1651 formed fewer filaments in higher concentrations of AMC (32/16 µg/ml) and IPM (16 µg/ml) corresponding to ½ MIC (**Table 1**). Upon reaching stationary phase growth in the presence of sub-inhibitory concentrations of antibiotics, β-lactam-induced cell filaments were no longer visible and morphology resembling cells unexposed to antibiotics was restored (**Fig. 1A**). Video imaging of Bp1651 grown in IPM below the MIC captures the dynamic nature of cell morphology over time (**Video 1**). In addition, the higher the drug concentration the longer cells remained elongated.

Visual observation of Bp1651 growing in the presence of sub-inhibitory concentrations of CAZ revealed increasing proportions of filamentous cells with increasing drug concentrations up to 64 µg/ml. To quantify this direct relationship, the lengths of 100 cells of strain Bp1651 were measured after 4 h in CAMHB alone and in broth containing two-fold increasing concentrations of CAZ. Histograms depict the distribution of cell lengths (**Fig. 1B**). In media without drug, 95% of cells measured ≤ 5 µm. For cells grown in broth with 8, 16, 32, or 64 µg/ml CAZ, the number of cells out of 100 with lengths ≥ 15 µm increased from 5, 16, 36, and 49 at each two-fold increasing concentration. The proportion of Bp1651 cells measuring ≤ 5 µm decreased substantially in 32 and 64 µg/ml, compared to 8 and 16 µg/ml CAZ, and cells reaching lengths of > 50 µm were observed at these concentrations (**Fig. 1B**). The median cell lengths for Bp1651 grown in CAMHB alone, and in broth containing 8, 16, 32, 64, and 128 µg/ml CAZ were 3, 4, 6, 11.5, 14 and 10.5 µm, respectively. The average cell lengths were slightly higher, 3.1, 5.6, 8.1, 15.1, 19.4, and 14.8 µm, respectively (**Fig. 1B**). The longest Bp1651 cell length (73 µm) was recorded in 64 µg/ml CAZ. More spheroplasts were visible after 6 h at the highest (128 µg/ml) CAZ concentration tested (**Fig. 1A**).

Growth kinetic graphs of Bp1651 in the presence of AMC and IPM show growth inhibition in CAMHB containing 64/32 µg/ml and 32 µg/ml, respectively (**Fig. S1A & Fig. S1B**). At these MICs, spheroplasts formed during exponential phase (**Fig. 1A**) and cell lysis began after 3.7 ± 0.0 h (AMC) and 3.6 ± 0.2 h (IPM) (**Table 1**). There was no evidence of cell filamentation at AMC or IPM MICs and median cell lengths were 4 and 3 µm, respectively.

### Cell morphology dynamics during *B. pseudomallei* β-lactam AST

Based on CLSI interpretative criteria for BMD testing for AMC, strain MSHR1655 is categorized as resistant (MIC of 32/16 µg/ml), 724644 is intermediate (MIC of 16/8 µg/ml), Bp6788 is susceptible with an MIC at the breakpoint (8/4 µg/ml) and 1026b is susceptible (MIC of 4/2 µg/ml). All *B. pseudomallei* study strains, except Bp1651, are CAZ-S and IPM-S based on conventional BMD testing (**Tables 1-3**). Optical microscopy was used to generate automated growth kinetic data and to acquire complementary video imaging of cell morphology dynamics of *B. pseudomallei* strains in the presence of AMC, CAZ, and IPM and in a broth only control (**Fig. 2** and **Fig. 3**). Morphology was monitored in antibiotic concentrations less than, equal to, and greater than MICs for each strain, which includes concentrations equivalent to the CLSI susceptibility breakpoints for each drug. All broth cultures were monitored over 18-20 h and optical screening images were captured during exponential phase growth after 6 h unless otherwise indicated. Average kinetic graphs (n=3) for each growth condition were overlaid on optical screen images and cell morphology dynamics are described below each image. In media without the addition of antibiotics, there was no evidence of cell elongation for *B. pseudomallei*, however, variable aggregation was observed between strains (**Fig. 2A**). Video imaging of strain 724644 showed cells amassing in groups during the first 10 h of growth in no-drug media (**Video 2**). Aggregation of cells was not observed for MDR strain Bp1651 or the susceptible Bp6788 in broth media alone (**Fig. 1A and Fig. S2**).

**Figure 2.**
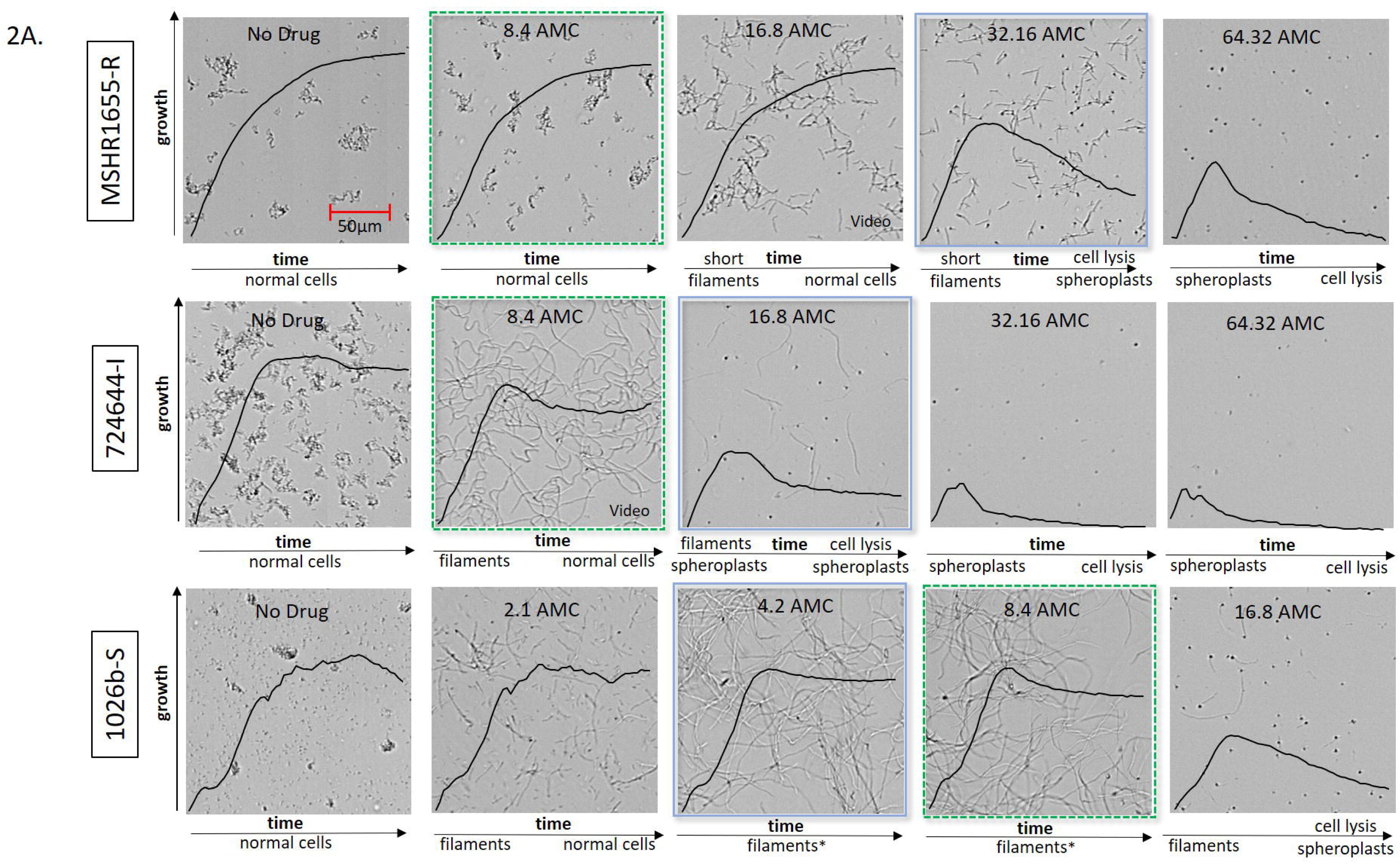
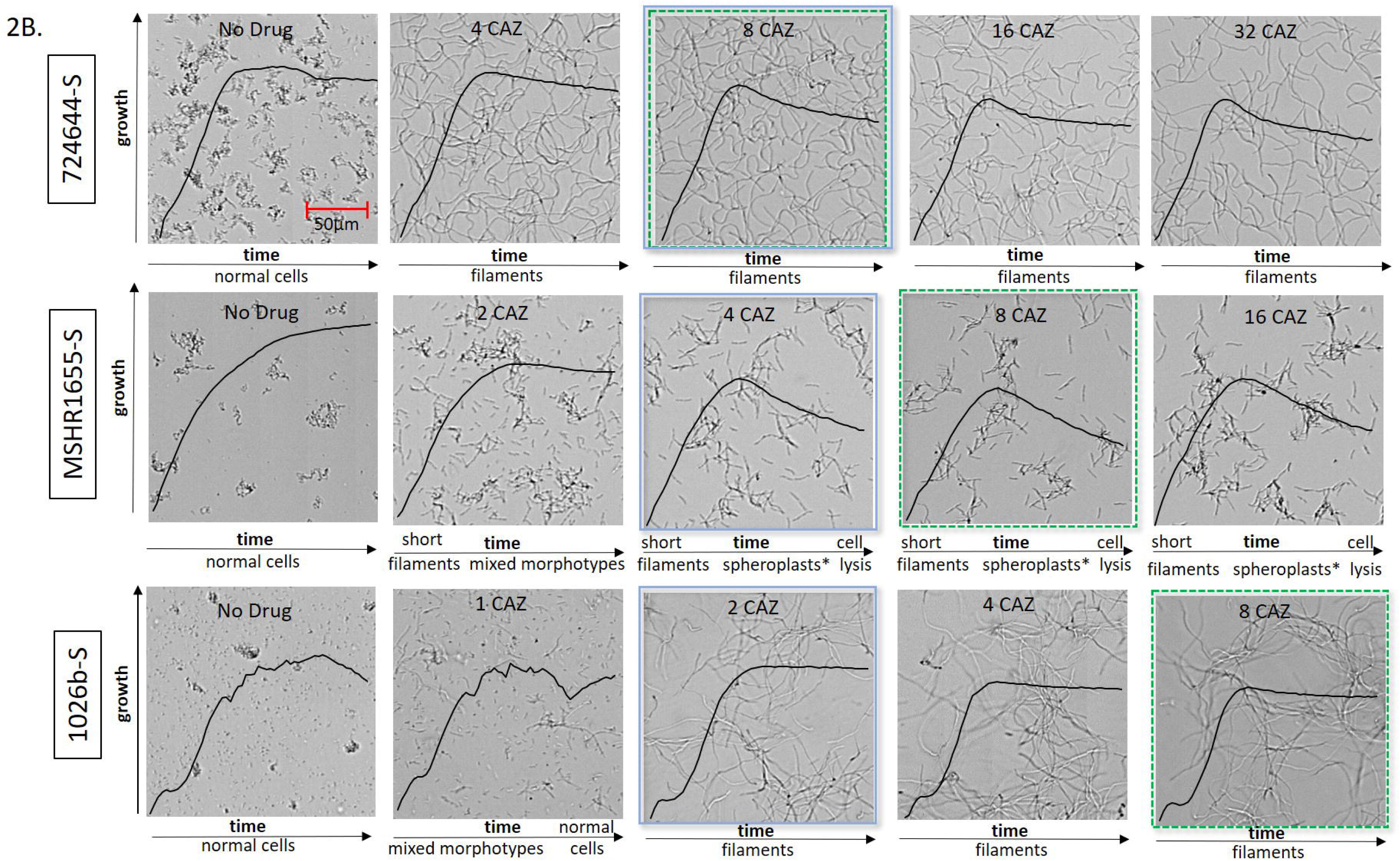
Cell morphology of *B. pseudomallei* strains cultured in the presence and absence of AMC (A) or CAZ (B). Optical screen images were captured after 6 h. Growth kinetic graphs (black lines) represent the average growth value of triplicate samples (y-axis) over time (18 h, x-axis) and morphology dynamics are described below each image. Strains are designated as resistant (R), intermediate resistant (I), or susceptible (S) to each antibiotic. Indication that a small proportion of spheroplasts were observed (*). Drug concentrations (µg/ml) tested were below, equal to, and above MICs (blue squares) for each strain. CLSI breakpoint for susceptibility (dotted green square), amoxicillin-clavulanic acid (AMC), and ceftazidime (CAZ).

**Figure 3.**
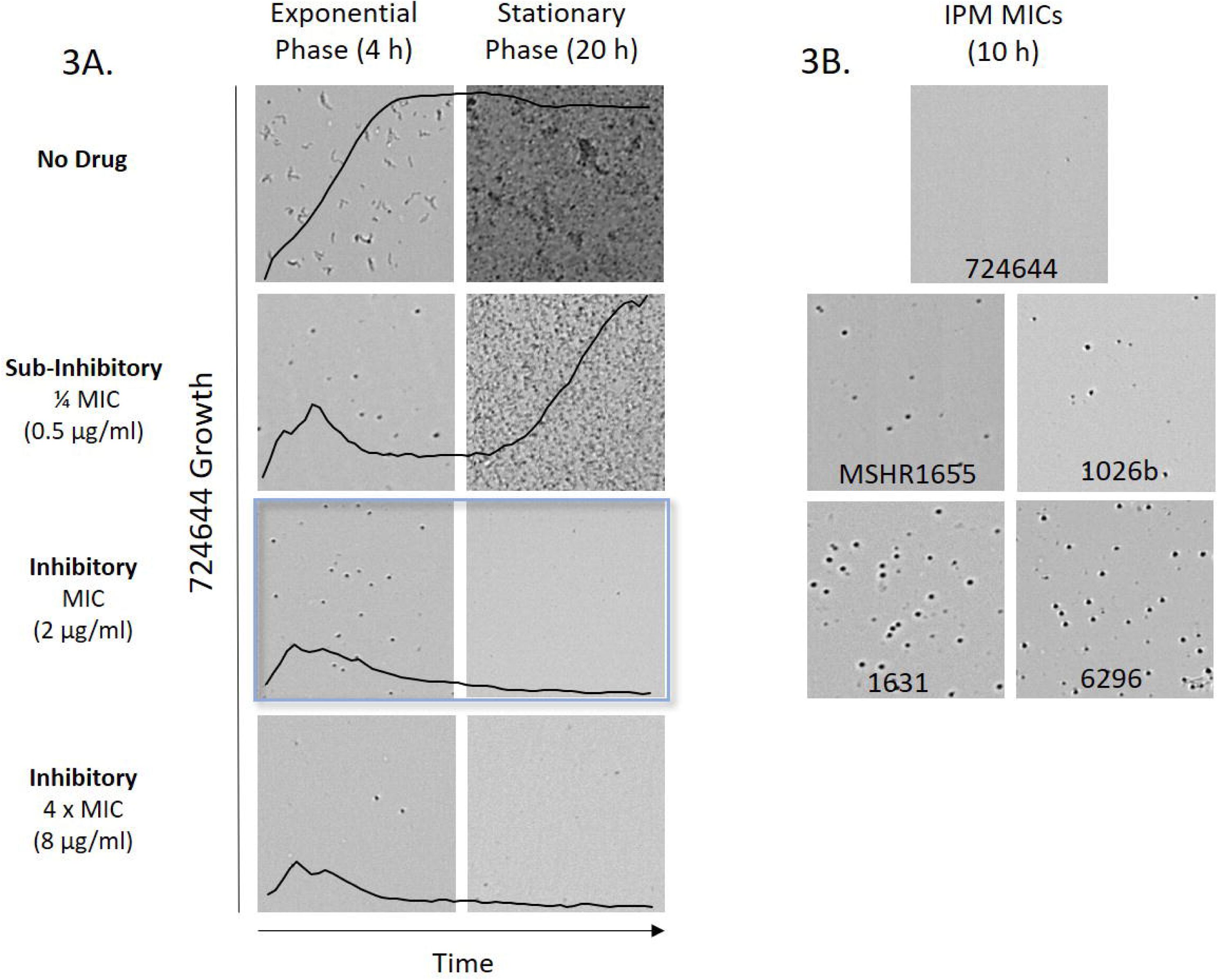
Cell morphology of *B. pseudomallei* strains cultured in the presence and absence of IPM. Optical screen images of 724644 (A) were captured during exponential phase growth (4 h) or during stationary phase (20 h). Imipenem (IPM) concentrations tested were below, equal to, and above the MIC (blue square). Growth kinetic graphs represent the average growth value of triplicate samples (y-axis) taken over time (20 h, x-axis). Optical screen images of *B. pseudomallei* strains in the presence of IPM MICs after 10 h (B).

### AMC

At sub-inhibitory concentrations of AMC, some filamentous cells were observed early on for the resistant, intermediate and susceptible *B. pseudomallei* strains (**Fig. 2A, Table 1 & 2**). At ½ MIC values, the number of cells ≥ 15 µm was variable between strains and ranged from 42/100 (1026b) to 93/100 (724644) (**Table 2**). The shortest filaments were observed for AMC-resistant strain MSHR1655. With the exception of Bp6788, during stationary phase growth in AMC concentrations below the MIC, replication of non-elongated cells was seen for the majority of strains. Video footage of strain 724644 grown in media containing AMC equivalent to ¼ (**Video 3**) and ½ (**Video 4**) the MICs demonstrates dynamic AMC-induced cell morphology changes over time. In the presence of 4/2 µg/ml AMC (or ¼ MIC), cells were slightly elongated over the first few hours of growth, with replication of cells resembling those not exposed to drug following quickly thereafter. At the CLSI breakpoint for susceptibility of 8/4 µg/ml AMC (or ½ MIC), 724644 cell filaments are considerably longer, and replication of non-elongated cells is detected much later in stationary phase (∼ 18 h).

Some evidence of cell filamentation was also observed for all strains during early exponential phase growth in broth with AMC at either the MIC or at 16/8 µg/ml, a concentration used to interpret *B. pseudomallei* susceptibility by conventional BMD (**Fig. 2A & Fig. S2**). Following initial filamentaion at MICs, formation of spheroplasts and cell lysis was observed for strains MSHR1655 and 724644 after 6.1 ± 0.2 and 4.8 ± 1.0 h, respectively (**Table 2**). Growth values initially increased in the growth kinetic graphs, and then subsequently decreased over time. However, detection of spheroplast formation and cell lysis could not be used to predict MICs as 1026b and Bp6788 remained filamentous over time at these AMC levels (**Fig. 2A**). Due to cell elongation, growth kinetic graphs could not accurately depict susceptibility, as microbial growth is based on changes in the surface area covered by all identified objects in a scan frame. Broth containing 4 x MIC was required to induce cell lysis of 1026b which commenced after 5.2 ± 0.4 h. Differences in cell morphology between strains early on at AMC MICs are apparent by the median cell length (MCL) values (n = 97-100) calculated after four hours which range from 6 to 44 µm (**Table 2**).

The length of 724644 cells was variable when measured in the presence of AMC below, at and above MICs for this strain. Cells of strain 724644 were measured in media without drug after two hours, without drug after four hours when cells had begun to aggregate, and in two-fold increasing concentrations of AMC after four hours (**Fig. 4a**). Distribution of cell lengths (µm) is depicted and MCLs are indicated by horizontal lines. Consistent with *B. pseudomallei* size, the MCL of strain 724644 grown in broth alone after two hours was 5.71 µm. Since the thinned length algorithm may not accurately identify overlapping cells as individual cells, the objects measured after four hours are more reflective of groups of aggregated cells, and at this time the MCL was 15 µm. At 8/4 µg/ml AMC, a sub-inhibitory concentration, the distribution of filament lengths was wide, with the longest cell reaching 126 µm (**Fig. 4A**). Unlike AMC-R strain Bp1651, in which only spheroplasts were detected at the MIC (64/32 µg/ml), both filaments and spheroplasts were observed at the MIC (16/8 µg/ml) for AMC-I strain 724644. Morphological heterogeneity of 724644 cells was documented throughout exponential phase prior to cell lysis (**Fig. 4B**). A heterogenous cell population, made up of both filaments and spheroplasts, was also observed for AMC-S strain Bp6788, but only in broth containing 2 x MIC (16/8 µg/ml) (**Fig. S2**). While the MCL was 6 µm, 15/100 cells were ≥ 20 µm. Increasing concentrations of AMC above the MICs induced the formation of more spheroplasts and resulted in quicker cell lysis for -R, -I and -S *B. pseudomallei* strains (**Fig. 2A & Table 2**)

**Figure 4.**
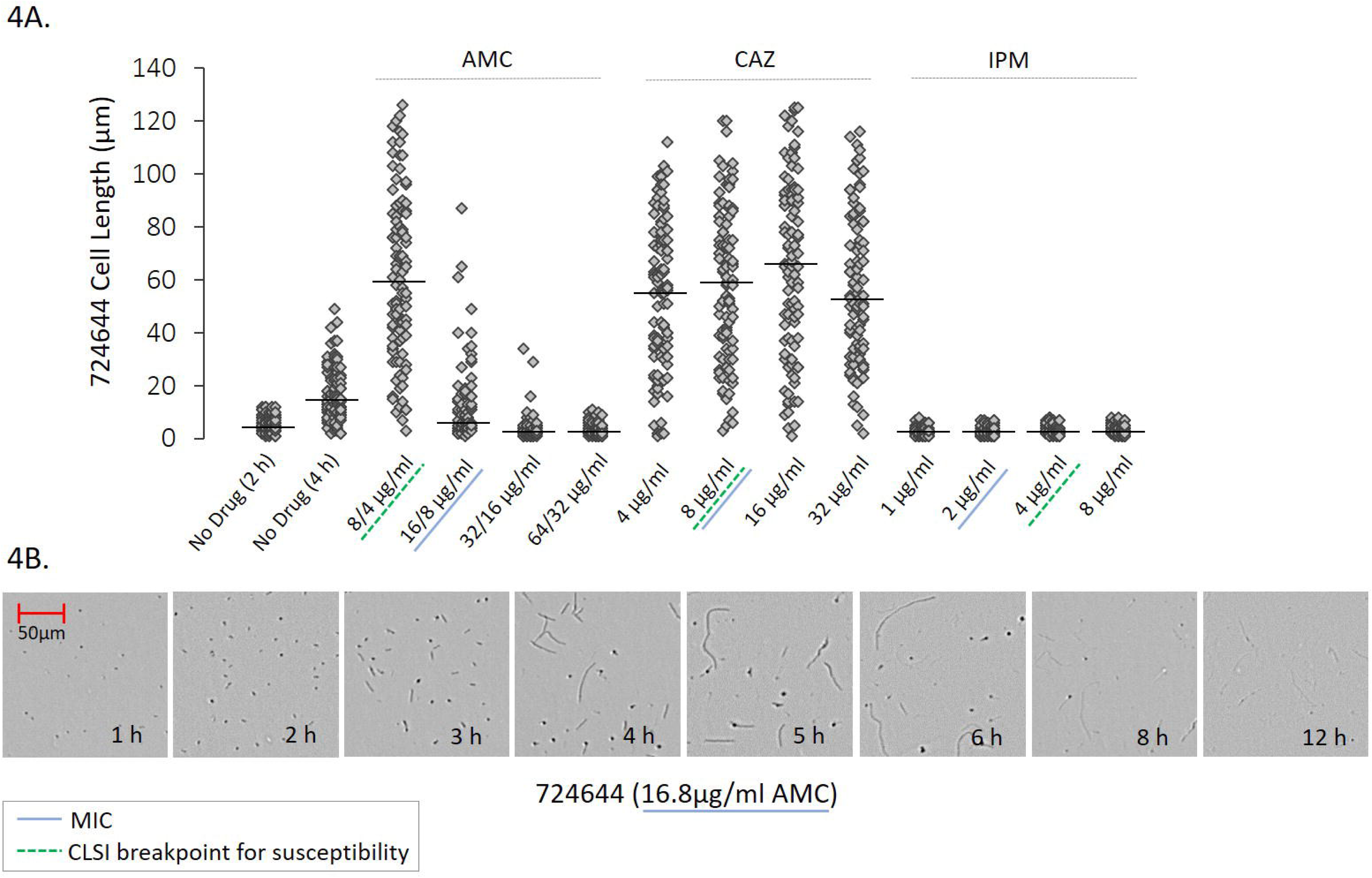
Cell length and morphology of strain 724644 cultured in the presence and absence of β-lactam antibiotics. The dot plot (A) represents the distribution of cell lengths (n=95-100) measured (µm) using the thinned length object feature. Cells in media without antibiotics were measured after 2 h and 4 h and cell exposed to amoxicillin-clavulanic acid (AMC), ceftazidime (CAZ) and imipenem (IPM) were measured after 4 h. Median cell lengths were calculated (black lines). CLSI breakpoint for susceptibility (green underline) and MICs (blue underline). Optical screen images taken over time (B) in the presence of the MIC value of AMC.

### CAZ

The CAZ MIC for strain 724644 was equivalent to the CLSI breakpoint for susceptibility (8 µg/ml). This strain formed long filaments in both sub-lethal and lethal concentrations of CAZ through all growth phases (**Fig. 2B**), with more than 90% of cells measured ≥ 15 µm after four hours (**Table 2**). Exposure to CAZ over time also induced long filaments for the susceptible 1026b in concentrations greater than or equal to the MIC, but not to the same degree in sub-lethal amounts at ½ MIC value (1 µg/ml) (**Fig. 2B**). As a result of filamentation, instrument-derived growth values could not be used to accurately determine susceptibility to CAZ. A wide distribution of cell lengths was noted for 724644 after four hours in all concentrations of CAZ tested, with MCLs between 53.5 and 66.5 µm (**Fig. 4**). Cells with lengths ≥ 100 µm were also recorded at each concentration. During exponential phase growth, CAZ induced the formation of shorter filaments for the susceptible MSHR1655 strain **(Fig. 2B**). In the presence of 4 and 8 µg/ml (MIC and 2 x MIC), MCLs were 22 µm. Unlike 724644, formation of spheroplasts and cell lysis was observed for MSHR1655 exposed to inhibitory CAZ concentrations, however these morphology changes did not occur rapidly. Cell lysis was observed after ∼ 8 h in concentrations of CAZ corresponding to the 1 to 4 x MIC (**Table 2**). Video imaging of MSHR1655 showed morphology changes including, formation of short filaments, followed by spheroplasts and subsequent cell lysis in media containing 16 µg/ml CAZ (**Video 5**). As growth values for MSHR1655 decreased in later time points due to cell lysis, susceptibility based on optical screening could be determined for this strain (**Fig. S1C**). At 8, 16, and 32 µg/ml CAZ, corresponding to the CLSI breakpoint and higher concentrations, susceptibility was determined after 9.2 ± 0.7, 9.3 ± 0.0, and 8.8 ± 0.2 h, respectively, with a 95% confidence level.

### IPM

Here, we performed rapid, optical screening-based AST and monitored IPM-induced morphology changes for five susceptible *B. pseudomallei* strains with MICs ranging from 0.25 to 2 µg/ml (**Table 3**). Unlike AMC and CAZ, cell filamentation was not observed in broth containing IPM at and above the MICs in video imaging of resistant or susceptible strains. Instrument-derived growth values could be used to accurately monitor bacterial replication in IPM. Since growth is inhibited for all susceptible *B. pseudomallei* strains at 8 µg/ml IPM based on CLSI interpretive criteria for conventional BMD, AST was performed at this concentration and at 16 µg/ml. Due to the rapid formation of spheroplasts followed by cell lysis induced by IPM for all *B. pseudomallei* strains tested, the time required to determine susceptibility was between 1.3 ± 0.9 and 3.8 ± 0.7 h in both concentrations (**Table 3**). To investigate whether cell lysis observations are useful to rapidly determine the strain MIC, the time in which this event occurred was also recorded for each *B. pseudomallei* strain. At these concentrations, time to cell lysis was variable between strains commencing as early as 3.0 ± 0.0 to h for 724644 and as late as 9.8 ± 0.2 h for Bp6296 (**Fig. 3A & Table 3**). Optical screening images of strains captured in the presence of MICs of IPM after 10 h showed more spheroplasts were present for strains with longer times to cell lysis (Bp6296 and Bp1631) (**Fig. 3B**). However, at 8 µg/ml IPM (2 to 32 x MIC values), the time range for cell lysis was narrower between strains, occurring between ∼ 3.0 to 6.0 h (**Table 3**).

In CAMHB containing IPM equivalent to ½ the MIC, cell filamentation was not observed during exponential phase growth for three of five study strains. Though, similar to IPM-R Bp1651, a small proportion of filamentous cells was seen in video footage of IPM-S strains Bp6296 and MSHR1655 during the first several hours. Some strains displayed an extended lag phase in sub-inhibitory levels of IPM. For instance, at ¼ MIC of IPM, after strain 724644 initially formed spheroplasts and underwent some cell lysis, a prolonged lag phase was observed until cells begin to replicate after 11 to 11.3 h (**Fig. 3A** and **Video 6**). No lag phase was observed for strain 724644 in lower concentrations of IPM, and rapid cell lysis was induced at and about the MIC (**Fig. 3A**). In media containing ½ IPM MIC, both resistant and susceptible strains Bp1651 (**Fig. S1B**) and Bp1631, respectively, displayed a lag phase lasting ∼ 9 h.

### *In silico* identification of penicillin binding proteins in *B. pseudomallei*

Ten genes encoding putative PBPs were identified in *B. pseudomallei* 1026b using the UniProtKB database. Based on conserved PBP domains predicted by Pfam and homology to PBPs in *P. aeruginosa* PAO1, strain 1026b contains five high molecular mass (HMM), multi-modular (containing both transglycosylase and transpeptidase domains), class-A PBP-1 homologs (*II0265, II0898, I3403, I1297*, and *II2482*). Four predicted HMM, class-B homologs containing transpeptidase domains were also identified; one PBP-2 (*I3332*) and three PBP-3s (*I0276, II1292* and *II1314*) as well as one low MM class-C PBP-6 (*I3098*) protein which contains a D-alanyl-D-alanine carboxypeptidase domain. NCBI Protein BLAST analyses revealed these *B. pseudomallei* 1026b PBP homologs share 29.2% – 45.5% identity to PBPs in *P. aeruginosa* PAO1. Ten corresponding PBP homologs were also encoded in the genomes of the resistant strains Bp1651 and MSHR1655 and the susceptible strain Bp6296.

## Discussion

Bacterial morphology is both complex and dynamic, and exposure to antibiotics can cause plasticity within a cell’s lifecycle. PBPs are essential for bacterial cell wall synthesis and the maintenance of cell shape. These proteins are also the targets for β-lactam antibiotics and their inactivation results in specific cell morphology changes. Numerous variables can affect β-lactam-induced morphological changes including the concentration, the duration of the exposure, the bacterial species, and the antimicrobial susceptibility [26-28]. PBP profiles and the binding affinity and kinetic interactions of β-lactams with PBPs are variable between species [29-31]. While PBPs and target affinities of β-lactam drugs are well described in *E. coli*, only genes encoding PBP-3 family homologs have been reported previously in *B. pseudomallei*, and PBP-1 and PBP-2 have not been documented. Here we identified 10 genes encoding putative PBPs in genomes of both susceptible and resistant strains, and these may represent the targets for β-lactams antibiotics. While the binding affinities of β-lactams for *B. pseudomallei* PBPs have not been investigated, drug-induced morphological changes can still lend insight into bacterial response and antibacterial mechanisms of action.

In *E. coli*, the primary affinity of amoxicillin is directed towards PBP-1s and PBP-2 [32]. The broad-spectrum β-lactamase inhibitor clavulanate binds to PBPs in both Gram-negative and -positive bacteria, including selectively to PBP-3 in *Streptococcus pneumoniae* [33]. Complementary binding to separate PBPs by a combination drug increased activity of single β-lactams used alone [33, 34]. In this work, we observed strain dependent cell morphology changes that varied based on AMC concentration and exposure time.

In sub-lethal concentrations of AMC, filament formation, to varying lengths and at variable proportions of cell populations, was evident for all *B. pseudomallei* strains tested. During exponential phase growth at AMC MICs, diverse cell morphologies were observed among strains including: formation of filaments, round spheroplasts, and a heterogenous population of both. Mixed morphologies in a cell population could indicate complementary binding to different PBPs. For all strains tested, higher concentrations of AMC at and above MIC values resulted in increased proportions of spheroplasts and/or subsequent cell lysis, which may indicate inactivation of PBP-2 and PBP-1. This is evident in the distribution patterns of strain 724644 cell lengths at higher AMC concentrations. The concentration of AMC that would ultimately lead to cell lysis was variable between strains and ranged from one to four times the MIC values for every *B. pseudomallei* strain evaluated. A change in cell morphology from filaments to spheroplasts or lysis has also been shown to increase with the duration of β-lactam exposure [35], which we observed herein. Morphological heterogeneity of cell populations was also observed in the presence of AMC for resistant and susceptible strains suggesting more than one PBP was inactivated.

At relatively low concentrations, CAZ was shown to bind to PBP-3 and caused rapid filamentation; at higher concentrations, cephalosporins can cause spheroplast formation or bind to PBP-1 resulting in cell lysis [28, 32, 36]. Here, at early time points susceptible *B. pseudomallei* strains 724644 and MSHR1655 formed long and short filaments, respectively, in CAZ concentrations below, at and up to eight times the MIC. Buijs *et al*. [26] studied CAZ-induced morphological changes of several Gram-negative species over 4 h and found filamentation started at sub or inhibitory concentrations and could extend far above MICs for *P. aeruginosa* and *Acinetobacter* spp. *Klebsiella* spp. formed shorter filaments, and one *K. pneumoniae* isolate displayed no filamentation in CAZ [26]. They concluded that PBP-binding specificities, which are reflected by morphology changes, differ for each genus or isolate. Based on our observations, CAZ may have a higher selective affinity for PBP-3 over time in strain 724644 which remained filamentous, whereas inhibition of a PBP-1 protein in MSHR1655 could be responsible for cell lysis at later time points. Previously, we showed CAZ-induced filamentation occurred at concentrations corresponding to 2 to 32 x MIC for susceptible *B. pseudomallei* and *B. mallei* strains, with cell lysis occurring in some strains after filament formation [20].

The resistant strain Bp1651 formed filaments early on in sub-inhibitory levels of CAZ. Then, original cell size, resembling those cells unexposed to antibiotics, was re-established at later time points. The time to morphology restoration was concentration dependent and may be attributed to the degradation of CAZ by the β-lactamase enzyme, PenA, which has acquired mutations correlating to high activity against this antibiotic [14]. In the presence of sub-inhibitory and inhibitory concentrations of cefotaxime, extended spectrum β-lactamase producing *E. coli* have also been shown to form filaments in lag and early exponential growth phases and revert back to non-elongated cells in exponential and stationary phase [37]. Chen *et al*. [38] demonstrated that filamentation of *B. pseudomallei* in sub-lethal CAZ concentrations could be reversed after antimicrobial removal and that reverted bacteria developed resistance. Other studies of *P. aeruginosa* showed cell morphology can be restored upon β-lactam removal [39].

For both susceptible and resistant *B. pseudomallei* strains, bacterial populations treated with CAZ resulted in a wide distribution of cell lengths. Bp1651 cells were shorter in the lowest and highest filament-inducing, sub-inhibitory CAZ concentrations. These observations are consistent with morphological studies of other Gram-negative species [26]. Median cell length of strain 724644 was more independent of CAZ concentration and cells with lengths greater than 100 µm were recorded in CAZ and AMC. Filaments with lengths up to 93 µm in the presence of sub- and lethal concentrations of β-lactams have been observed [40].

In the presence of IPM at ½ the MIC, a small proportion of filamentous cells was observed in the first few hours for the resistant strain Bp1651 and for two of five susceptible *B. pseudomallei* strains. Exposure to meropenem (MEM) for two hours was shown to induce filamentation in both carbapenem-resistant *P. aeruginosa* mutants and their susceptible parents at this concentration [41]. At the IPM MIC, *B. pseudomallei* strains formed spheroplasts at an early growth phase, similar to *A. baumannii* [42]. These cells are morphologically typical of spheroplasts, with a near-perfect spherical shape and found as individuals rather than in cell arrangements like ovoid cells [43]. Future studies may assess the osmotic stability of these round cells, as spheroplasts are characteristically instable [28, 44].

Satta *et al*. [45] defined the relationship between cell killing kinetics and PBP binding in *E. coli*, by demonstrating IPM concentration-dependent PBP inhibition. Saturation of more than one PBP resulted in different rates of cell lysis. Here, at IPM MICs (0.25 to 2 µg/ml), the exposure time resulting in bacteriolysis was variable among *B. pseudomallei* isolates (∼ 3 to 10 h). At 8 µg/ml, an IPM concentration used to interpret *B. pseudomallei* susceptibility by conventional BMD, the exposure time for induction of cell lysis narrowed (∼ 3 to 6 h). It is plausible that the rate of PBP-1 and PBP-2 saturation is variable between strains and may explain the different times to bacteriolysis.

After spheroplast formation and some cell lysis was observed at ½ and ¼ the IPM MIC for strains Bp1651 and 724644, respectively, morphology and exponential phase growth were restored at later time points. In clinical isolates of *E. cloacae* and *Klebsiella* spp., carbapenem tolerance was mediated by cell wall-deficient spheroplasts [46]. It is suggested that Gram-negative pathogens have the ability to survive for extended periods without structurally sound cell walls and that morphological recovery and cell division are possible upon removal of antimicrobials [46]. β-lactam-treated isolates from patients were shown to contain spheroplasts, and the high tolerance observed for some clinical isolates cannot solely be explained by rare persistor cells alone [46-48]. Recurrence of *B. pseudomallei* infection is one of the most relevant complications in melioidosis survivors and treatment and prophylaxis must be tailored to individual patients according to clinical manifestations and response [49, 50]. Investigations studying the survival of *B. pseudomallei* spheroplasts exposed to and then removed from the presence of IPM/MEM *in vitro* and *in vivo* may inform therapeutic decision making, especially if carbapenem levels drop or if antibiotic administration is discontinued.

Novel approaches, including high-powered optical microscopy, microfluidic assays, flow cytometry, and dielectrophoresis systems, allow for cellular-level observation of bacterial morphologies [20, 21, 23, 51]. These analyses have been used to develop rapid antimicrobial susceptibility tests for several Gram-negative organisms. Su *et al*. [23] used a dielectrophoretic system to accurately and rapidly determine susceptibility of several Gram-negative spp. to cefazolin, CAZ, cefepime, and doripenem based on morphology changes. Within 90 min at MICs, they showed β-lactams induced cell shape changes in susceptible strains such as elongation, cell swelling or cell lysis; however, cell morphology remained unchanged for resistant strains [23]. Here, at MICs and at concentrations that would be used to interpret AMC and CAZ susceptibility by conventional BMD, both resistant and susceptible *B. pseudomallei* strains demonstrated morphology changes early on. Filamentation could not be used as rapid indicator of susceptibility. Evidence of elongation was observed for resistant strains, and cell lysis could not determine an MIC since filamentation was observed for several strains at these concentrations. Even in IPM, the time to cell lysis was variable between strains and did not always occur rapidly.

Otero *et al*. [22] also developed a rapid (75 min) assay to detect antimicrobial resistance of Gram-negative spp. (*A. baumanni, K. pneumonia*, and *P. aeruginosa*) based on cell elongation in CAZ and established concentration ranges in which susceptible strains increased in length, but resistant strains did not. These drug ranges were several dilutions lower than the CLSI breakpoints used for conventional BMD susceptibility testing for these pathogens. While we performed β-lactam AST using concentrations below, equal to, and above MICs, including at those that could be used to interpret susceptibility by BMD, it is possible that morphological differences between susceptible and resistant *B. pseudomallei* strains may be assessed more accurately at lower concentrations. Choi *et al*. [21], developed a rapid AST based on single-cell morphological analysis and reported that some resistant Gram-negative bacteria, such as *P. aeruginosa*, can both deform in shape and still divide in the presence of IPM and piperacillin; this observation is accounted for in their assay. Recent development of a microplate-based surface area assay for rapid phenotypic AST also allows for more accurate measurements of replication when bacteria filament or swell [52].

Due to the complexity and dynamic nature of bacterial morphology, developing a rapid β-lactam AST based on cell shape alone proves complicated. We demonstrate *B. pseudomallei* morphology is dependent on strain, susceptibility profile, β-lactam exposure time and antibiotic concentration. Quantitative morphological data represents a snap shot in time of continuously fluctuating bacterial cells. Observations of morphological heterogeneity of certain cell populations in the presence of β-lactams highlight the need for a large number of bacteria to be tested to avoid generalizations of response. We also showed, in sub-inhibitory concentrations, resistant strains undergo filamentation during early growth phase, similar to susceptible strains. Despite the limited number of resistant *B. pseudomallei* strains available in our collection to evaluate and the biosafety and biosecurity challenges associated with working with this pathogen, trends in morphology could be used to inform both bacterial response to β-lactams and antibiotic mechanisms of action. Identification of putative PBPs in the *B. pseudomallei* genome reveals the possible targets for β-lactams. Future work characterizing these PBPs and binding affinities of β-lactams may build on this foundation. A greater understanding of β-lactam-induced cell morphology changes could contribute to more meaningful clinical decisions for melioidosis patients or provide critical strain-specific information during an outbreak or public health emergency.

## Conclusions

Using optical microscopy, we describe the morphology dynamics of *B. pseudomallei* strains with distinct AST profiles, exposed to clinically relevant β-lactam antibiotics. Ten genes encoding putative PBPs in the *B. pseudomallei* genome were identified and represent potential targets for β-lactams. Both resistant and susceptible strains exhibited filamentation during early exposure to AMC and CAZ at concentrations used to interpret susceptibility (based on CLSI guidelines). While developing a rapid β-lactam AST based on cell-shape alone requires more extensive analyses, growth attributes of *B. pseudomallei* reveal information about antibiotic response and antibacterial mechanisms of action.

## List of Abbreviations

(AST): Antimicrobial susceptibility test,
(BMD): broth microdilution,
(CLSI): Clinical and Laboratory Standards Institute,
(IPM): imipenem,
(CAZ): ceftazidime,
(AMC): amoxicillin-clavulanic acid,
(MDR): multidrug-resistant,
(R): resistant,
(I): intermediate,
(S): susceptible,
(SBA): Trypticase Soy Agar II with 5% sheep blood,
(CAMHB): cation-adjusted Mueller Hinton broth,
(MIC): minimal inhibitory concentrations,
(TES): N-tris(hydroxymethyl) methyl-2-aminoethanesulfonic acid,
(SESA): Segmentation and Extraction Surface Area,
(SD): standard deviation,
(PBPs): penicillin-binding proteins.

## Declarations

## Ethics approval and consent to participate

Not Applicable.

## Consent for publication

Not Applicable.

## Availability of data and material

Datasets used and analyzed for this study are available from the corresponding author upon request.

## Authors Contributions

Design and conception of experiments: HM, DS. Execution of experiments: HM and JB. Data analysis: HM. Drafting the manuscript: HM, DS. Corrections and approval of the final manuscript: HM, JB and DS. All authors read and approved the final manuscript.

## Acknowledgements

We thank Jay E. Gee in the Division of High-Consequence Pathogens and Pathology at the Centers for Disease Control and Prevention for his review of this manuscript. We also acknowledge the team at BioSense Solutions ApS (Farum, Denmark) for their technical support.

## Disclaimer

The findings and conclusions in this report are those of the authors and do not necessarily represent the official position of the Centers for Disease Control and Prevention. Use of trade names is for identification only and does not imply endorsement by the Centers for Disease Control and Prevention.

## Supplemental Figures

**Figure S1. Growth kinetics of *B. pseudomallei* strains evaluated over 18 h in the presence and absence of** β**-lactams.** Bp1651 exposed to AMC (A) and IPM (B), MSHR1655 exposed to CAZ (C) and 724644 exposed to IPM (D). Drug concentrations (µg/ml) corresponding to the CLSI breakpoint for susceptibility (green dots) and MICs (blue dots). Graphs represent the average ± standard deviation from triplicate samples. Amoxicillin-clavulanic acid (AMC), ceftazidime (CAZ), and imipenem (IPM).

**Figure S2. Cell morphology of *B. pseudomallei* 6788 in the presence and absence of AMC (µg/ml).** Optical screen images were captured after 6 h. Amoxicillin-clavulanic acid (AMC), CLSI breakpoints for susceptibility (green dotted line), MIC (blue line).

