## Supplementary figures and images for "Optical microscopy reveals the dynamic nature of *B. pseudomallei* morphology during β-lactam antimicrobial susceptibility testing"

### Fig S1

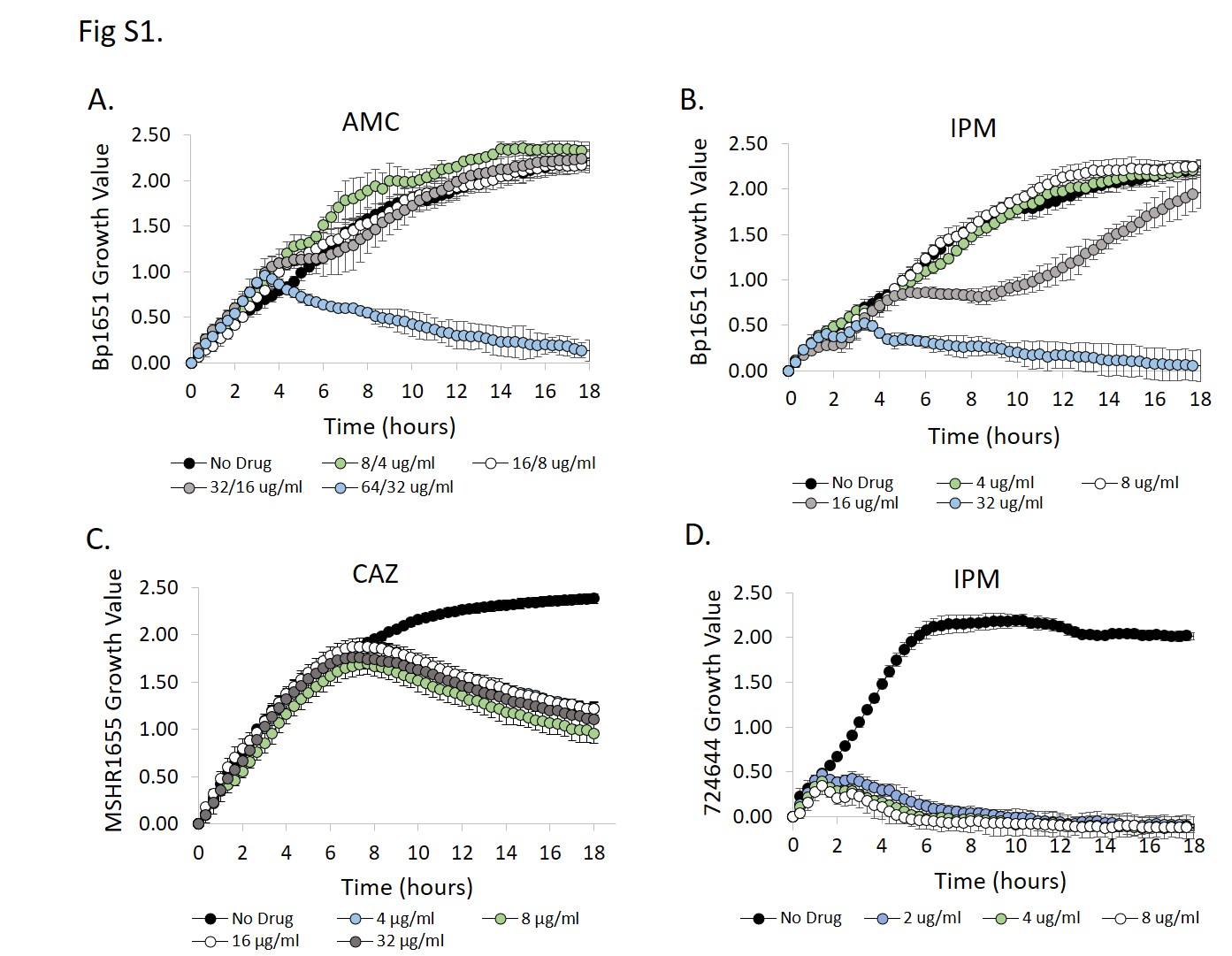

### Fig S2

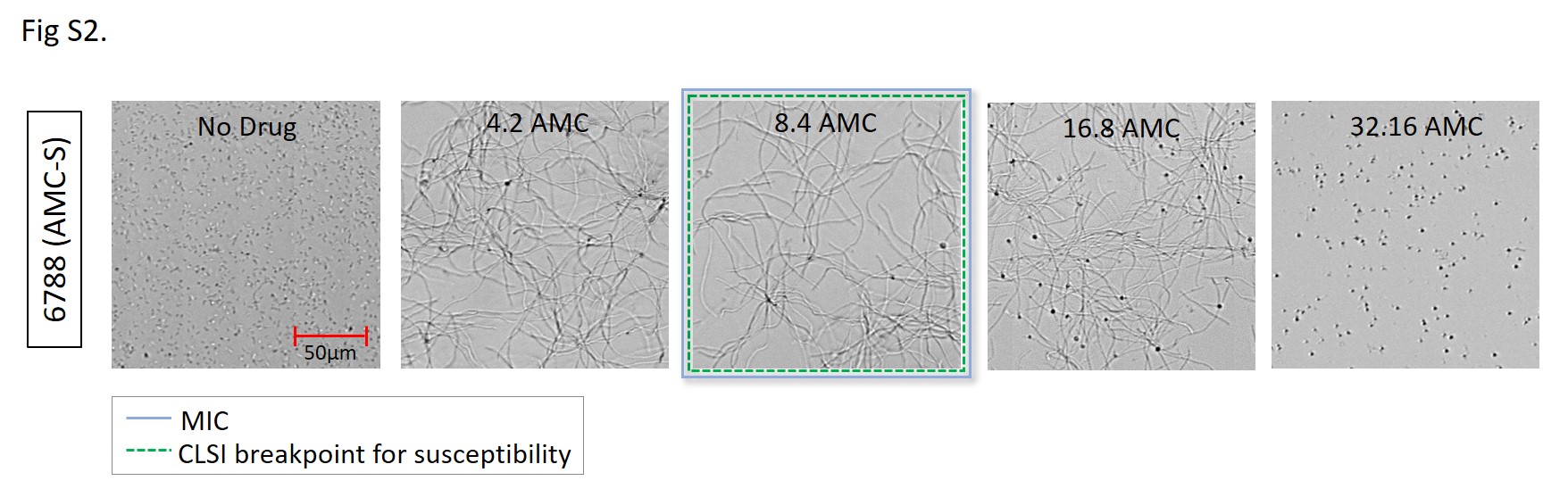
